# SARS-CoV-2 type I Interferon modulation by nonstructural proteins 1 and 2

**DOI:** 10.1101/2022.06.09.495586

**Authors:** Émile Lacasse, Isabelle Dubuc, Leslie Gudimard, Annie Gravel, Isabelle Allaeys, Éric Boilard, Louis Flamand

## Abstract

Since the beginning of the COVID-19 pandemic, enormous efforts were devoted to understanding how SARS-CoV-2 escapes the antiviral response. Yet, modulation of type I interferons (IFNs) by this virus is not completely understood. Using *in vitro* and *in vivo* approaches, we have characterized the type I IFN response during SARS-CoV-2 infection as well as immune evasion mechanisms. The transcriptional and translational expression of IFNs, cytokines and chemokines were measured in lung homogenates of Wuhan-like, Beta, and Delta SARS-CoV-2 K18-ACE2 transgenic mice. Using *in vitro* experiments, we measured SARS-CoV-2 and its non-structural proteins 1 and 2 (Nsp1-2) to modulate expression of IFNβ and interferon-stimulated genes (ISG). Our data show that infection of mice with Wuhan-like virus induces robust expression of *Ifna* and *Ifnb1* mRNA and limited type I production. In contrast, Beta and Delta variant infected mice failed to activate and produce IFNα. Using *in vitro* systems, *Ifnβ* gene translation inhibition was observed using an Nsp1 expression vector. Conversely, SARS-CoV-2 and its variants induce robust expression of NF-κB-driven genes such as those encoding CCL2 ans CXCL10 chemokines. We also identified Nsp2 as an activator of NF-κB that partially counteracts the inhibitory actions of Nsp1. In summary, our work indicates that SARS-CoV-2 skews the antiviral response in favor of an NF-κB-driven inflammatory response, a hallmark of acute COVID-19, and that Nsp2 is partly responsible for this effect.

**Importance:** Several studies suggest that SARS-CoV-2 possess multiple mechanisms aimed shunting the type I interferon response. However, few studies have studied type I IFN modulation in the context of infection. Our work indicates that mice and human cells infected with SARS-CoV-2 produce sufficient type I IFN to activate an antiviral response, despite Nsp1 translational blockade of *IFNΒ1* mRNA. In contrast to Wuhan-like virus, Beta and Delta variants failed to induce *Ifna* gene expression. Our work also showcases the importance of studying protein functions in the context of infection, as demonstrated by the partial antagonizing properties of the Nsp2 protein on the activities of Nsp1. Our studies also highlight that the innate immune response triggered by SARS-CoV-2 is chiefly driven by NF-κB responsive genes for which Nsp2 is partially responsible.

## Introduction

Severe acute respiratory syndrome coronavirus 2 (SARS-CoV-2) emerged in Wuhan, China, in December 2019 leading to the coronavirus infectious disease 2019 (COVID-19) global outbreak (1). SARS-CoV-2 is a member of the *Coronaviridae* family, *Orthocoronavirinae* subfamily, *Betacoronaviruses* genus, *Sarbecovirus* subgenus (2). SARS-CoV-2 genome consists of a single-stranded (ssRNA) positive RNA genome of an approximate length of 29,7 kb (2, 3). The SARS-CoV-2 genome encodes 4 structural proteins (spike (S), envelope (E), membrane (M), and nucleocapsid (N)), 7 accessory proteins (ORF3a, ORF6, ORF7a, ORF7b, ORF8, and ORF10) and ORF1ab, a large open reading frame (ORF) which encodes a large polyprotein which gets cleaved in 16 non-structural proteins (Nsp1-16) (2). SARS-CoV-2 infection implicates the binding of the S protein to the human Angiotensin-Converting Enzyme 2 (ACE2) followed by the cleavage of the S2 subunit by transmembrane protease serine protease-2 (TMPRSS-2) and ADAM metallopeptidase domain 17 (ADAM17) (4, 5). Finally, SARS-CoV-2 enters its host cell by endocytose (6).

One of the first host defense mechanisms against pathogens like viruses is the innate immune system that is initiated by the recognition of pathogen-associated molecular patterns (PAMPs) by cellular sensors (7). One of the main systems triggered in response to viral infections is interferon (IFN) production (8). In the case of infection by viruses like SARS-CoV-2, the type I IFN pathways can be activated by two different processes. One of them involves the recognition of double-stranded RNA (dsRNA), produced during SARS-CoV-2 replication, by RIG-I like receptors (RLRs) such as retinoic acid-inducible gene I (RIG-I) and/or melanoma differentiation gene 5 (MDA5) sensors located in the cytoplasm (9, 10). Viral RNA recognition by these sensors leads to the phosphorylation, dimerization, and nuclear translocation of IFN regulatory factor 3 (IRF3) and IRF7. In parallel, NF-κB activation is initiated and together with IRF3/7, *IFNB1* gene transcription is initiated (11–13). Type I IFN transcription can also be activated by the recognition of dsRNA by the Toll-like receptor 3 (TLR3) or by the recognition of ssRNA by TLR7/8 (14). TLR activation results in *IFNβ* gene transcription through similar signaling pathways (15).

The products of type I IFN genes, IFNα/β1, are secreted in the extracellular space. IFN receptor (IFNAR1-2) engagement activates the Janus kinases signal transducer and activator of transcription proteins (JAK-STAT) pathway that leads to the expression of many dozen interferon-stimulated genes (ISGs) whose products are responsible for establishing the antiviral defense (16–18).

In addition to activating IFN signaling, recognition of viral PAMPs, as well as damage-associated molecular patterns (DAMPs) generated by viral replication, by TLRs, NOD-like Receptors (NLRs) or AIM2 like receptors (ALR), lead to the activation of NF-κB targeted gene and the formation of the inflammasome. These pathways can initiate the production and activation of several pro-inflammatory mediators such as cytokines and chemokines (19, 20).

Without surprise, various components of the type I IFN response are targeted by many viruses and other pathogens. As was observed with SARS-CoV, some viral proteins, such as Nsp1, can target signaling proteins and modulate the immune response of the host (21, 22). Furthermore, recent studies indicate that SARS-CoV-2 Nsp1 can evade the type I IFN response by inducing translational shutdown (23, 24). Conversely, SARS-CoV-2 Nsp2 seems to amplify the type I IFN response (23, 25, 26). Most studies conducted on type I IFN response evasion by SARS-CoV-2 were carried out using single protein expression systems that cannot fully recapitulate infection or conditions where several viral and cellular proteins are expressed simultaneously. On the opposite, some studies suggest that SARS-CoV-2 induces type I IFN expression (27). In the current study, we have used K18-hACE2 mice infected with different strains of SARS-CoV-2 and a human pulmonary epithelial cell line to characterize the IFN response during infection. We also examined the effects of Nsp2 on the ability of Nsp1 to shut down IFN synthesis. Our results suggest that all three SARS-CoV-2 isolates modulate IFNβ1 similarly while the Beta and Delta variants are much more effective in preventing IFNα production than the original Wuhan-like strain. Moreover, our work suggests that the translational shutdown mediated by Nsp1 is the main mechanism capable of inhibiting IFNβ1 production and that Nsp2 dampens this inhibitory activity, in part through the activation of the NF-κB pathway. Our results argue that SARS-CoV-2 skews the antiviral response in favor of an NF-κB driven inflammatory response and highlights the caveat of studying viral proteins outside the context of infection.

## Materials and Methods

### Cell culture and virus

HEK293T and Vero cells were purchased from American Type Culture Collection (Manassas, VA, USA), A549-hACE2 and HEK293T-hACE2 cells were obtained from Biodefense and Emerging Infections Research Resources Repository (BEI Resources, Manassas, VA, USA). These cell lines were passaged twice a week. HEK293T and A549 cells were cultured in Dulbecco’s Modified Eagle Medium (DMEM) (Corning Cellgro, Manassas, VA, USA) with 10 % fetal bovine serum (FBS) (Corning Cellgro), 10mM HEPES pH 7.2, 1% (v/v) nonessential amino acid (Multicell Wisent Inc., St-Bruno, QC, Canada) and 5μg/mL of *Plasmocin*® (Invivogen, San Diego, CA, USA), to prevent mycoplasma contamination. Vero cells were cultured in Medium 199 (Multicell Wisent Inc.) supplemented with 10 % FBS and 5μg/mL of *Plasmocin*®. Cell lines were grown at 37°C with 5% CO_2_. Sendai virus (SeV) was obtained from Charles River Laboratory (Saint-Constant, QC, Canada) and SARS-CoV-2 Wuhan-like strain (LSPQ, B1 lineage) from the Laboratoire de Santé Publique du Québec ([LSPQ] Sainte-Anne-de-Bellevue, QC, Canada), this strain will be considered as a wild-type strain. SARS-CoV-2 Beta strain was obtained from BEI resources and SARS-CoV-2 Delta strain from the BC CDC. SARS-CoV-2 strains were propagated on Vero cells and the supernatant of infected cells was used for infection experiments. The infectious titer of the Wuhan-like strain viral preparations was 1.8 x10^6^ Tissue Culture Infectious Dose 50/mL (TCID_50/mL_) for mice experiments and 5.24 x10^6^ TCID_50/mL_ for *in vitro* experiment, 1.80×10^6^ TCID50_/mL_ for Beta strain and 2.08×10^6^ TCID_50/mL_ for Delta strain. A549-hACE2 were infected with Wuhan-like strain at a multiplicity of infection (MOI) of 1 for one hour, cells were then washed 2 times with Phosphate-buffered saline 1X (PBS) and new culture media was added. Experiments involving infectious SARS-CoV-2 viruses were performed in a BSL-3 facility.

### Determination of the viral titer

Vero cells were plated in a 96-well plate (2×10^4^/well) and infected with 200µl of serial dilution of the viral preparation or lung homogenate in the M199 media supplemented with 10mM HEPES pH 7.2, 1mM of sodium pyruvate, 2.5g/L of glucose, 5μg/mL *Plasmocin*® and 2% FBS. Three days post-infection plates were analyzed using a EVOS M5000 microscope (ThermoFisher Scientific, Waltham, MA, USA) and the viral titer was determined using the Kerber method.

### Mice

B6.Cg-Tg(K18-hACE2)2Prlmn/J (stock#3034860) mice were purchased from the Jackson Laboratories (Bar Harbor, ME). Nine-week-old male and female mice were infected with 25μL of saline containing 9×10^3^ (TCID_50/mL_) of the corresponding SARS-CoV-2 strain or 25μL of saline for mock-infected mice. Mouse weight was recorded every day until euthanasia. Mice were sacrificed on day 3 post-infection and lungs were collected for RNA extraction and tissue homogenization for cytokines and infectious titer (TCID_50/mL)_ analysis.

### Plasmids and reagents

SARS-CoV-2 non-structural protein (Nsp) 1 and 2 expression vectors were generated by amplifying the genes from SARS-CoV-2 RNA. Nsp1 and 2 genes were cloned into pENTR (L1-L2) using Hifi DNA Assembly (New England Biolabs, Ipswich, MA, USA). LR recombination Gateway (ThermoFisher Scientific) was used to recombine Nsp1 and Nsp2 genes into pCDNA5-TO (obtained from Dr. Anne Claude Gingras, Lunenfeld-Tanenbaum Research Institute, Toronto, On, Canada). To generate an Nsp1-Nsp2 polyprotein coding vector, Nsp1-P2A (28) was cloned into the pcDNA5-TO-Nsp2 vector using PCR overlap cloning with Hifi DNA Assembly. Expression vector IFN-β-LUC was obtained from Dr. Nathalie Grandvaux (CHUM, Montreal, QC, Canada), ISRE-LUC (*Interferon-sensitive response element*) expression vector was obtained from BD Biosciences (Mississauga, ON, Canada), the NF-κB-LUC expression vector was obtained from Michel J. Tremblay (CHUL, Quebec, Qc, Canada), PRD1-III-LUC vector was obtained from Dr. Tom Maniatis (Zuckerman Institute, Colombia, USA). Polyinosinic-polycytidylic acid (poly(I:C)) was purchased from Cytiva (Mississauga, ON, Canada). BCA Protein Assay kit was purchased from (ThermoFisher Scientific). Primers used for plasmid construction are listed in the supplementary table 1.

### Transfection

HEK293T cells were transfected using TransIT®-LT1 (Mirus, Madison, WI) reagent with indicated expression vectors. Poly(I:C) transfections were done using lipofectamine 3000 reagent (ThermoFisher Scientific) at a ratio of 1:5.

### Protein expression

HEK293T cells were plated in a 6-well plate (6.5×10^5^/well) 24h before transfection. A549-hACE2 cells cultured in a 6-well plate (2×10^5^ cells) were infected with SARS-CoV-2. Forty-eight hours post-transfection, cells were lysed in radio-immunoprecipitation assay buffer (RIPA) buffer with HALT protease Inhibitor Cocktail (ThermoFisher Scientific) or directly in Laemmli 2X buffer. The proteins were separated by SDS/PAGE gel and transferred to a PVDF low fluorescence membrane (Bio-Rad Laboratories Ltd, Mississauga, ON, Canada). Membranes were incubated with 1*μ*g/mL of mouse anti-Flag (Applied Biological Materials Inc., Richmond, BC, Canada), 0.25 *μ*g/mL of rabbit anti-SARS-CoV-2-Nsp1 (Genetex, Irvine, CA, USA), or 0.25 *μ*g/mL of rabbit anti-SARS-CoV-2-Nsp2 (Genetex) for 1 hour at room temperature or 16 hours at 4°C. Peroxidase-labeled goat anti-mouse IgG (Jackson Immunoresearch Laboratories Inc., West Grove, PA, USA) (40 ng/mL) or peroxidase-labeled goat anti-rabbit (Jackson Immunoresearch Laboratories Inc.) (80 ng/mL) were used as secondary antibodies for 1 hour at the room temperature and revealed with the addition of Clarity Western ECL reagent (Bio-Rad Laboratories Ltd). ChemiDoc MP Imaging System (Bio-Rad Laboratories Ltd) or radiological films (Mandel, Guelph, ON, Canada) were used to capture images. Rabbit anti-tubulin 2A and 2B (Abcam Inc., Toronto, ON, Canada) (0.66μg/mL) or mouse anti-tubulin (ThermoFisher Scientific) (0.33μg/mL) or Stain-Free Imaging Technology® (Bio-Rad Laboratories Ltd) were used as loading controls.

### Reporter assays

HEK293T cells were plated in 24-well plates (1.6 x10^5^/well) and transfected with 50ng to 100ng of reporter vectors and 30ng to 300ng of Nsp1, Nsp2 or Nsp1-Nsp2 vectors brought to 0.5μg/well with the empty expression vector. Twenty-four-hour post-transfection, transfected cells were infected with 20 hemagglutinin units of SeV or stimulated with 500 units of IFNα (PBL Assay Science, Piscataway, NJ, USA). Sixteen hours later, cells were lysed and the luciferase activity was determined as previously described (29)

### IFNß induction and ISG induction

HEK293T-hACE2 cells were plated (3×10^4^/well) in 12-well plates and transfected with 0.2μg of Nsp1 vector, 0.6μg of Nsp2 vector, or 0.8μg of Nsp1-Nsp2 vector complete to 1μg/well with empty pCDNA5. Twenty-four-hour post-transfection, cells were infected with 40 hemagglutinin units of SeV for sixteen hours. Supernatants and cells were collected separately, and cells were lysed in 0.5mL of QIAzol reagent (Qiagen, Toronto, ON, Canada). Samples were stored at −80°C until future analysis.

### Infection and poly(I:C) stimulation

A549-hACE2 cells were plated (7.5 x10^4^/well) in 12-well plates and infected with Wuhan-like SARS-CoV-2 strain following the same procedure described above. Twenty-four hours post-infection, cells were transfected with 2μg/mL of poly(I:C) for 16 hours. Supernatants and cells were collected separately, and cells were lysed in 0.5mL of QIAzol reagent (Qiagen). Supernatants were incubated with 1% triton for one hour at room temperature to inactivate SARS-CoV-2. Samples were stored at −80°C until analyzed.

### IFNß quantification

IFNβ in the supernatant was quantified with the Human IFN-beta DuoSet enzyme-linked immunosorbent assay (ELISA) kit, according to the supplier recommendations (R&D Systems Inc., Toronto, ON, Canada).

### Multiplex cytokines quantification

Cytokines in mouse lung homogenates were measured using a custom ProcartaPlex^TM^ Mouse Mix & Match Panels kit (Invitrogen Waltham, MA, USA) on the Bio-Plex 200 (Bio-Rad Laboratories Ltd).

### Quantitative real-time PCR analysis

Total RNA from cell cultures was extracted following QIAzol protocol and RNA from mouse lungs was extracted using the Bead Mill Tissue RNA Purification Kit and the Omni Bead Ruptor Bead Mill homogenizer (Kennesaw, GA). Following extraction, residual DNA was removed by treating the samples with DNAse I (Roche, Mississauga, ON, Canada). For the quantification of human gene expression and mouse *Cxcl1*, *Ccl2*, *Isg56*(*Ifit1*), *Ifny*, and *Ifna*, RNA was reverse transcribed to cDNA using SuperScript™ IV VILO™ mastermix (ThermoFisher Scientific). Quantitative real-time PCR (qPCR) w performed using the SsoAdvanced Universal Probes Supermix (Bio-Rad Laboratories Ltd) for *IFNB1* gene and *GAPDH* as the housekeeping gene. SsoAdvanced Universal SYBR Green Supermix (Bio-Rad Laboratories Ltd) was used for human *ISG15* and *ISG56* and mouse genes including *Gapdh* as the housekeeping gene on the Rotor-Gene Q 5plex (Qiagen). RT-qPCR primers and probes are listed in the supplementary table 2.

### Digital PCR analysis

SARS-CoV-2 viral RNA loads were determined using Droplet Digital PCR (ddPCR) supermix for probes without dUTP (Bio-Rad Laboratories Ltd) and the QX200 Droplet Digital PCR System Workflow (Bio-Rad Laboratories Ltd). ddPCR primers and probes are listed in the supplementary table 2.

### RT^2^ profiler PCR Arrays

RNA extracted from mouse lungs as described above was cleaned up using On-Column DNAse using RNAse-Free DNase Set (Qiagen) and RNeasy Mini Kit (Qiagen). RNA was reverse transcribed using RT^2^ First Strand Kit (Qiagen). qPCR and quality control were done using RT^2^ SYBR^®^ Green ROC FAST Mastermix (Qiagen) and RT^2^ profiler PCR Arrays: Mouse Antiviral response (Qiagen). Data analyses were performed using the GeneGlobe (Qiagen) analyzing tool. Genes of the RT2 profiler PCR Arrays are listed in the supplementary table 3.

### Immunofluorescence

A549-hACE2 cells were plated (1.6×10^4^/well) in 8-well chamber slides. Twenty-four hours later, cells were infected with SARS-CoV-2 as described above. Forty-eight hours post-infection, cells were fixed in 2% paraformaldehyde in PBS for one hour at room temperature. Cells were then incubated for thirty minutes in blocking solution (PBS with 0.1% bovine serum albumin [BSA], 3% FBS, 0.1% Triton X-100 and 1mM EDTA) then with 17μg/mL of rabbit anti-SARS-CoV-2-N (Rockland Immunochemicals Inc., Limerick, PA, USA) in the blocking solution for one hour at room temperature. After, cells were washed three times for 5 minutes with PBS and incubated with 4μg/mL of goat anti-rabbit-Alexa-488 (ThermoFisher Scientific) in the blocking solution for 30 minutes at room temperature. Finally, cells were washed for 5 minutes in PBS and incubated into PBS 1X with 1.67ug/mL of DAPI (Invivogen). Cells were washed for 5 minutes in PBS, mounted with ProLong Diamond Antifade reagent (ThermoFisher Scientific) and images were acquired using a Z2 confocal microscope with LSM 800 scanning system (Zeiss, Germany). Images were captured with a 20x objective (Zeiss, Apochromat). ZEN 2.3 software (Zeiss, Germany) was used to acquire and process images. Z-stack projections of 3 μm in total thickness are represented.

## Results

### Viral loads and host gene modulation following infection by Wuhan, Beta, and Delta viruses in K18-hACE2 mice

To model the innate immune response against the Wuhan, Beta, and Delta SARS-CoV-2 strain, we infected K18-hACE2 mice with a lethal dose of virus. Three days post-infection, cytokine load and antiviral related genes were measured in mouse lungs. No change in body weight or temperature was observed at this time (30). As shown in Fig. 1A, the copy number of the SARS-CoV-2 E gene/genomic RNA was almost threefold higher in Beta-infected mice relative to Wuhan-infected mice. Viral RNA loads for Delta-infected mice were higher than Wuhan-infected mice, but this difference was not statistically significant. When the infectious viral loads from the same lung homogenates were analyzed, all three groups of mice showed similar infectious viral loads (Fig. 1B). The difference between these two viral quantification methods is since viral RNA quantification measure gRNA and mRNA while infectious viral loads measure only infectious viral particles.

**Figure 1.**
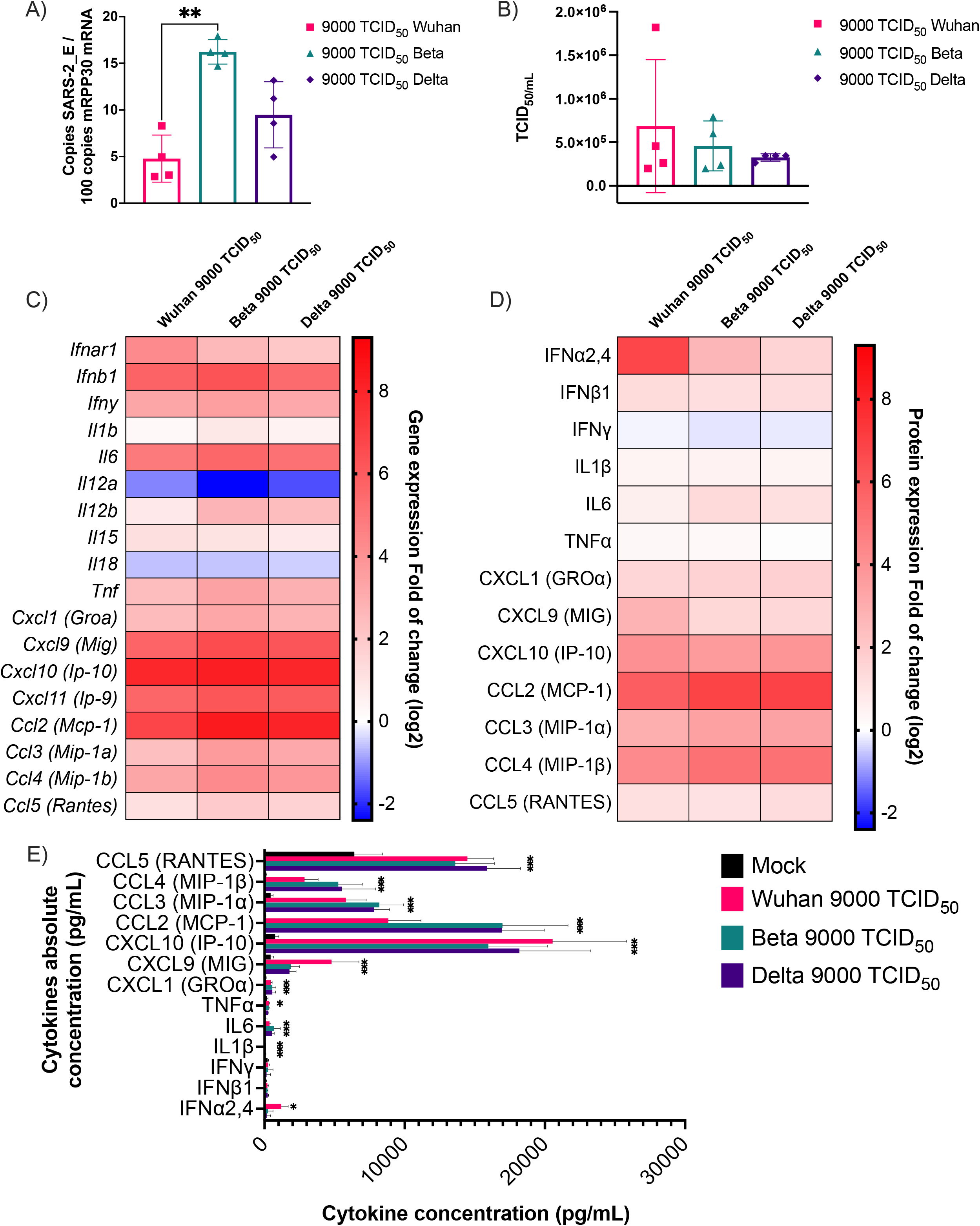
Cytokine mRNA and protein expression profile following infection of K18-ACE2 mice with Wuhan, Beta and Delta strains. Infected or mock mouse lung tissues were collected three days post-infection (n=4/group). A) The number of SARS-CoV-2 *E* gene copy number was evaluated by ddPCR using lung RNA and expressed as copie number per 100 copies of *Rpp30* mRNA. (B) Infectious viral titers were determined in lung homogenates and expressed in TCID_50/mL_. (C-D) Gene expression was evaluated by RT-qPCR and cytokine concentration in lung homogenates determined using a 13-plex Luminex panel. Cytokine gene expression and concentration levels are presented as heatmaps with results expressed as fold (log_2_) relative to mock-infected mice. Statistical analyses were done by comparing 2^(- ΔCt)^ values for each gene in the control group and infected groups with a nonparametric T-test and only data with a p value less than 0,05 are show. (E) Absolute cytokine concentrations in lung homogenates. Results are expressed as mean +/-SD (n=4 mice/group). For protein quantification, statistical analyses were done by comparing the normalized concentration for each cytokine in the control group and infected groups with a nonparametric T-test. *P<0,05, **P<0,01, ***P<0,001, ****P<0,0001.

### SARS-CoV-2 and its variants induce a robust chemokine production but a limited type I IFN production in mice

Chemokine C-X-C Ligand 10 (*Cxcl10* [*Ip-10*]), Chemokine C-C Ligand 2 (*Ccl2* [*Mcp-1*]), *Cxcl11 (Ip-9), Cxcl9* (*Mig*), *Ifnb1* and Interleukin 6 (*Il-6*) were the main cytokine genes upregulated by all variants (Fig. 1C). *Ifny*, Tumor necrosis factor (*Tnf*)*, Cxcl1*(*Groa*)*, Ifna, Ccl3* (*Mip-1a*), *Ccl4* (*Mip-1b)* genes were also upregulated, but to a lesser extent. On the opposite, *Il11a* and *Il18* genes were downregulated. When analyzed at the protein level, CC and CXC chemokines were efficiently produced in response to infection. In contrast, despite robust *Ifnb1*, *Ifny*, *Il6*, and *Tnf* gene expression, little gene products were measured (Fig. 1D). Of potential interest, the Wuhan strain induced the gene expression and the release of IFNα while Beta and Delta variants did not. Despite, the IFNα and IFNβ1 protein production, chemokine production was one thousand to twenty thousand time higher (Fig. IE) than pro-inflammatory cytokines and type I IFN production. These results show that SARS-CoV-2 innate immune response in mice was dominated by chemokines.

### SARS-CoV-2 infection in mice does not induce an inflammatory reaction mediated by the inflammasome

As well as the cytokines mentioned above, the expression of several genes involved in the Toll-Like receptors (TLRs), NOD-Like receptors (NLRs) and RIG-Like receptors (RLR) signaling pathways were measured. Despite upregulation of the Mediterranean fever gene (*Mefv*) implicated in the inflammasome formation and proinflammatory cytokine release (31), inflammasome components such as Apoptosis-associated speck-like protein containing a CARD (*Pycard*), Proline-serine-threonine phosphatase-interacting protein 1(*Pspip1*), Absent in melanoma 2 (*Aim2*), Caspase 1 (*Casp1*) and NLR family pyrin domain containing 3 (*Nlrp3*) were not modulated early in infection (Fig. 2A). Moreover, Caspase recruitment domain-containing protein 9 (*Card9*) and Mitogen-activated protein kinase 14 (*Mapk14*), implicated in the inflammatory process, were downregulated. This finding is consistent with the lack of proinflammatory cytokines such as *Il1b* and *Il18* (Fig. 1C).

**Figure 2.**
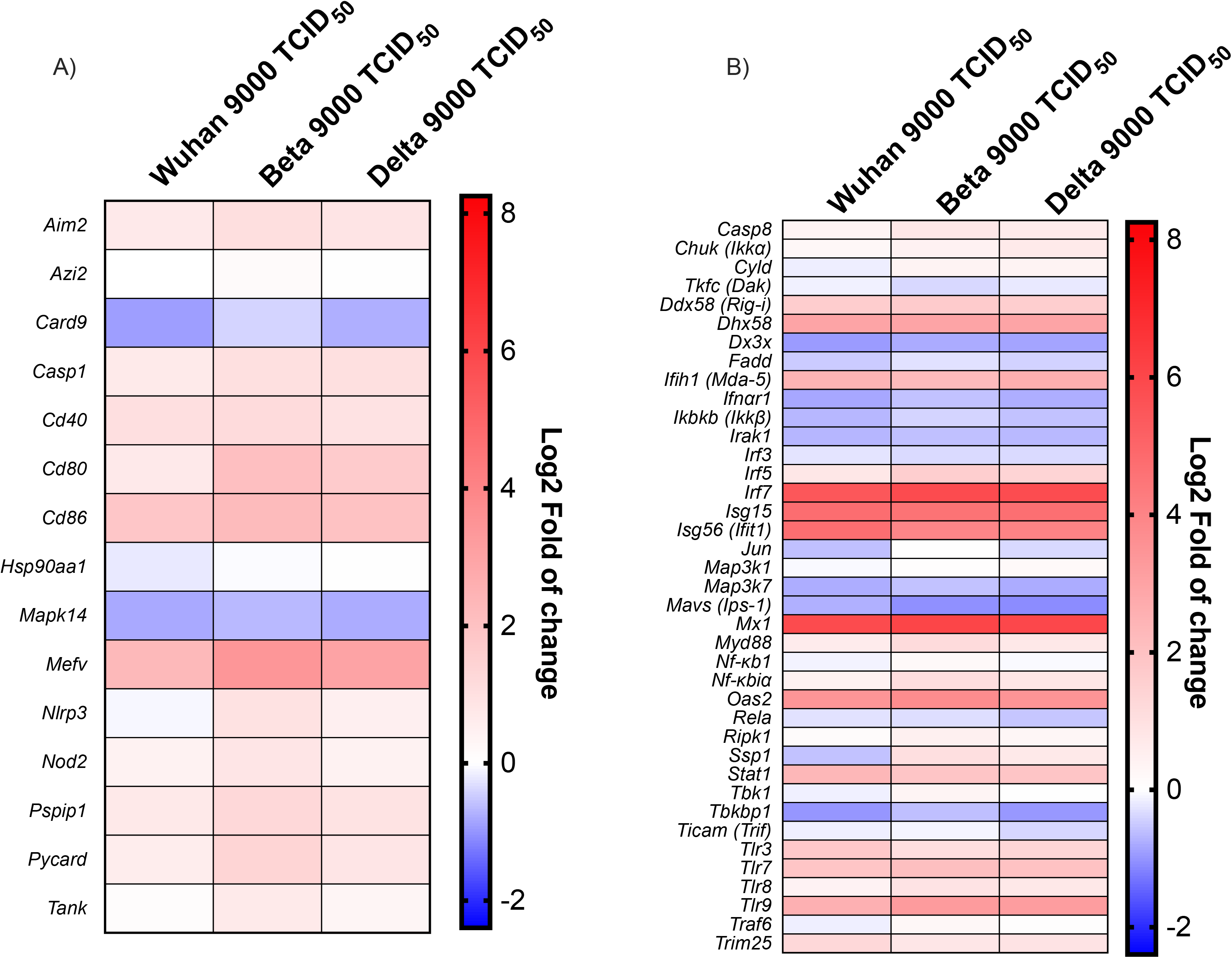
Antiviral response gene expression following infection with Wuhan, Beta and Delta strains. Heat map representation of cytokines and inflammatory related genes (A) and Type I IFN production and signalisation related genes (B). Results are expressed as fold (log_2_) relative to mock-infected mice. For gene expression, statistical analyses were done by comparing 2^(- ΔCt)^ values for each gene in the control group and infected groups with a nonparametric T-test and only data with a p values less than 0,05 were show. For protein expression, statistical analyses were done by comparing normalised concentration of each cytokine in the mock infected groups with infected groups with a nonparametric T-test. *P<0,05, **P<0,01, ***P<0,001, ****P<0,0001.

### Modulation of the IFN activation pathways during SARS-CoV-2 infection

As shown in the Fig. 2B, many downstream effector genes such as Inhibitor of nuclear factor kappa-B kinase subunit beta (*Ikbkb*[*Ikkb*]), Interleukin-1 receptor-associated kinase 1 (*Irak1*), Transcription factor (*Jun*), *Mavs* [*Ips-1*], Mitogen-activated protein kinase kinase kinase 7 (*Map3k7*[*Tak1*]) were downregulated during infection. Moreover, Canopy FGF signaling regulator 3 (*Cnpy3*), a TLRs chaperon (32), was also downregulated which could impair the recognition of viral PAMPs. On the other hand, cytoplasmic and endosomal ssRNA and dsRNA sensors, such as *Tlr3, Tlr7, Tlr8*, DExD/H-box helicase 58 (*Ddx58*[*Rig-i*)), 2’-5’-oligoadénylate synthetase 2 (*Oas2*) and Interferon-induced helicase C domain-containing protein 1 (*Ifih1* [*Mda-5*]), were upregulated during SARS-CoV-2 infection (Fig. 2B). Moreover, *Irf7* gene transcription was also robustly induced following the infection by each viral strain (Fig. 2B).

### Robust ISGs expression despite low-level type I IFN production during SARS-CoV-2 infection

The expression of genes associated with type I IFN signaling was monitored during infection (Fig. 2B). ISGs such as *Isg15*, *Isg56* (IFIT1), and Interferon-induced GTP-binding protein Mx1 (*Mx1*) were strongly induced following infection with all three SARS-CoV-2 strains. In agreement with those results, *Stat1* gene transcription was also upregulated by the different infections. On the opposite, *Ifnar1* expression was downregulated.

### SARS-CoV-2 infection induces *IFNβ* gene transcription, inhibits IFNβ protein synthesis, and does not affect type I IFN signaling

A549-hACE2 were infected with the SARS-CoV-2 Wuhan strain and the efficiency of infection was visualized by immunofluorescence using anti-nucleocapsid (N) antibodies (Fig. 3A). IFNβ mRNA was quantified by RT-qPCR in mock-infected or SARS-CoV-2-infected cells with or without poly(I:C) stimulation, a type I IFN inducer. IFNβ mRNA quantification indicated that SARS-CoV-2 efficiently activated *IFNB1* gene transcription. Stimulation with poly(I:C) amplified *IFNB1* mRNA expression (Fig. 3B). Similar results were obtained for *ISG15* and *ISG56* gene expression +/-poly(I:C) (Fig. 3C and D). Conversely, no IFNβ1 protein production was detected in the supernatant of infected cells with SARS-CoV-2 and the infection partially inhibiting the poly(I:C) mediated activation (Fig. 3E).

**Figure 3.**
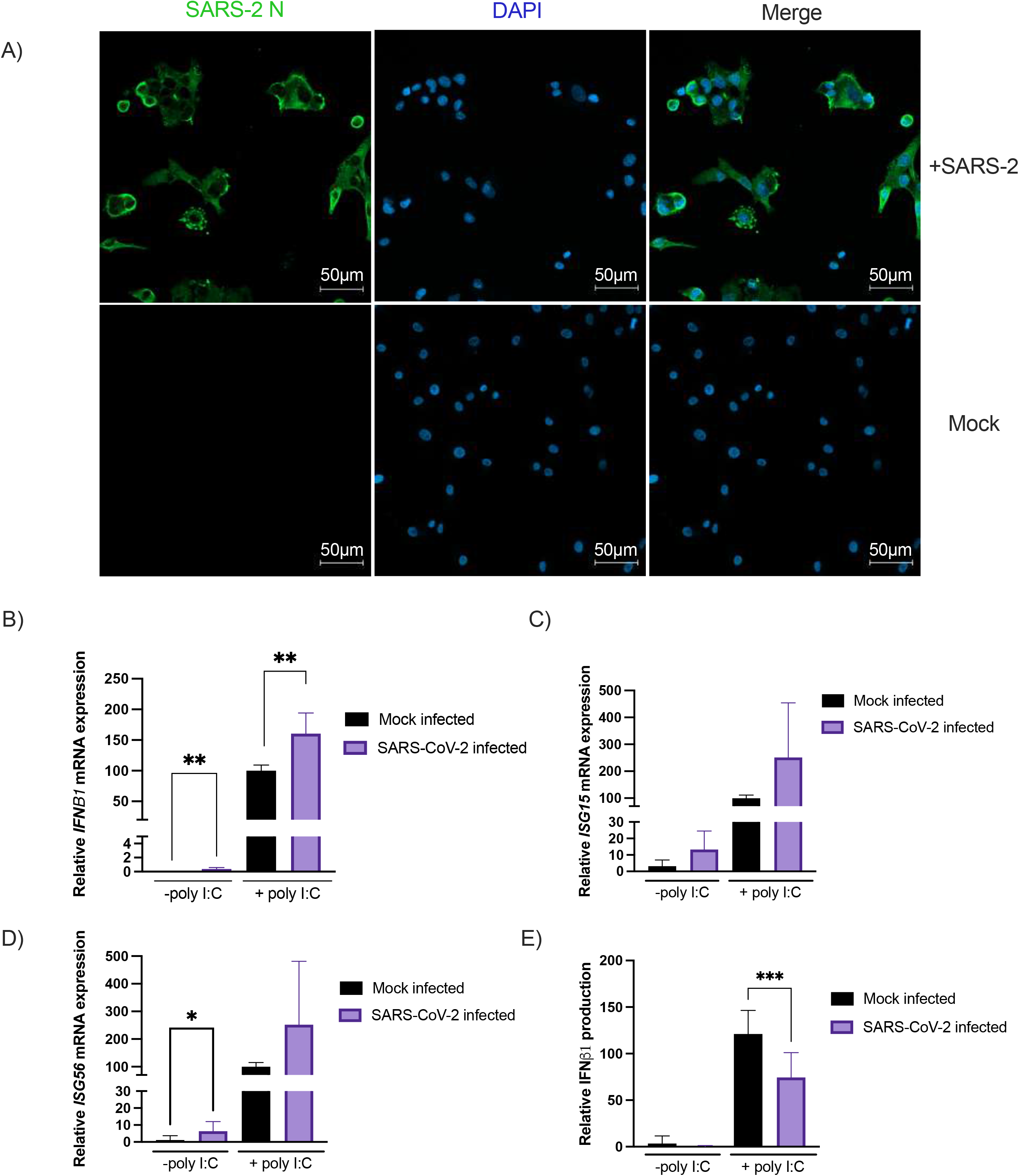
SARS-CoV-2 Nucleocapsid staining on A549-hACE2 infected cells. Forty-eight hours post-infection cells were fixed and stained as described in the materials and methods section (A). Effect of SARS-CoV-2 infection on poly(I:C) *IFNB1* mRNA expression (B), *ISG15* mRNA (C) or *ISG56* mRNA expression (D), induced IFNβ1 production (E). 24h post-seeding, A549-hACE2 cells were infected with SARS-CoV-2 as described in the materials and methods section. 32h post-infection, SARS-CoV-2 and mock infected cells were stimulated with poly(I:C) and RNA extracted and analyzed with RT-qPCR while IFNβ1 was measured in the supernatant by ELISA. All experiments were performed twice in triplicate and the compilation of the data is shown. Error bars indicate SD of biological triplicate repetitions. Statistical analyses were done by comparing mock control with corresponding condition with a nonparametric T-test. *P<0,05, **P<0,01, ***P<0,001, ****P<0,0001.

### Effects of SARS-CoV-2 Nsp1 and Nsp2 on IFNβ1 and type I IFN responsive promoters

Despite expressing elevated levels of IFNβ1 mRNA, SARS-CoV-2-infected cells synthesize a limited amount of IFNβ1 protein, suggesting that viral factors affect mRNA translation. Work by others (23, 24) has indicated that the Nsp1 protein is a potent mRNA translation inhibitor. During infection, Nsp1 is generated by the proteolytic cleavage of a large precursor protein yielding several additional proteins, including Nsp2. Thus, we studied Nsp1’s behavior in the absence of Nsp2. Nsp1 and Nsp2 expression vectors were co-transfected into HEK293T cells with IFNβ promoter or the ISRE promoter luciferase reporters. As shown in Fig. 4A and C, Nsp1 strongly inhibited the SeV-induced IFNβ1 promoter activation and the IFNα-induced ISRE activation in a dose-dependent manner. On the opposite, Nsp2 expression activated IFNβ1 and ISRE promoters and amplified the responses to SeV and IFNα (Fig. 4 B and D). To determine whether the expression levels of Nsp1 and Nsp2 derived from expression vectors were physiologically relevant, these were compared to Nsp1 and Nsp2 levels measured during infection. As shown in Fig. 4E and F, Nsp1 and Nsp2 relative protein expression in SARS-CoV-2 infected cells was in the expression range of transfection doses used for reporter assays.

**Figure 4.**
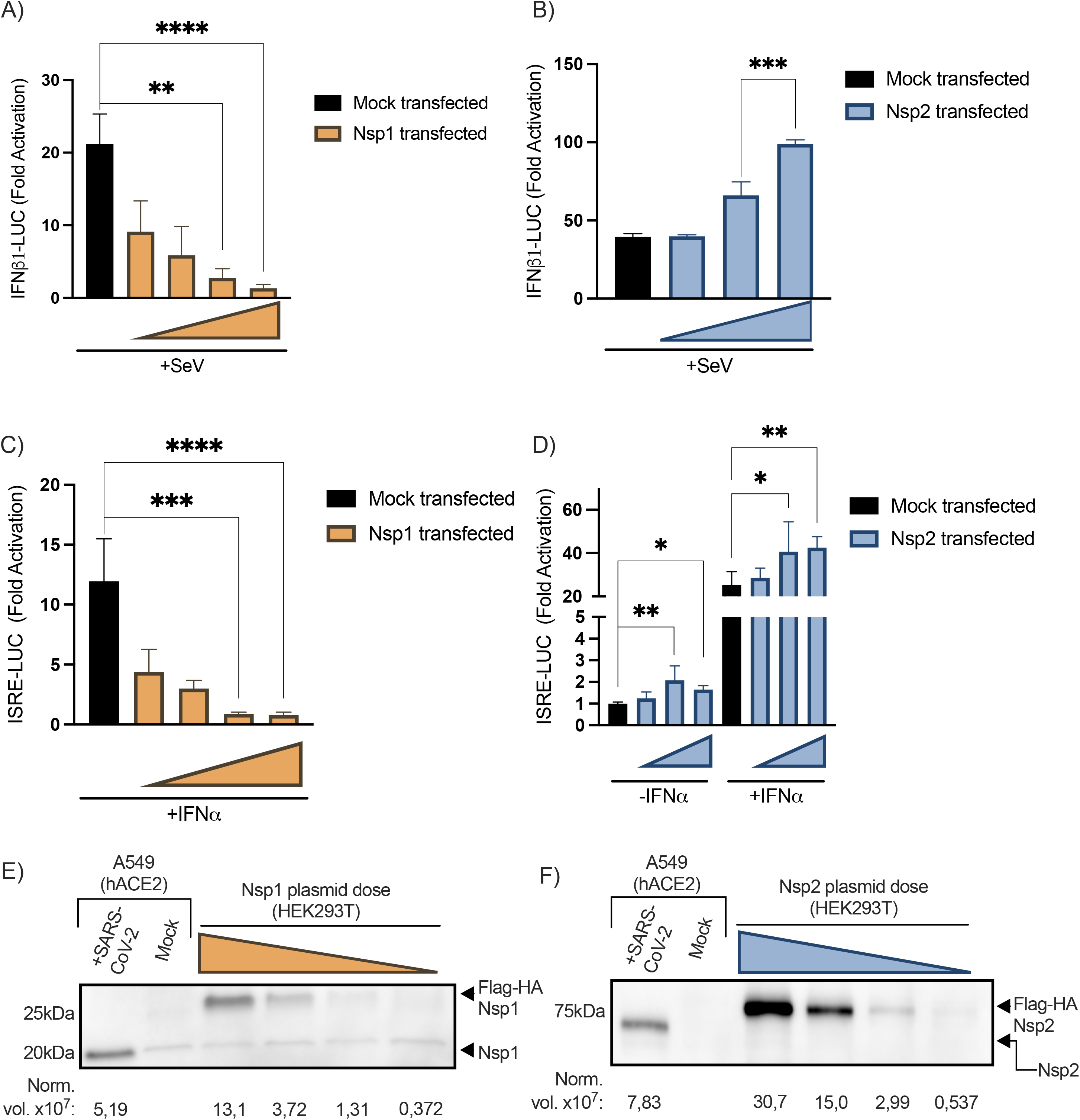
Effect of SARS-CoV-2-Nsp1 or Nsp2 on IFNβ1-luc and ISRE-luc promoter activation. 24h post-seeding, cells were transfected with IFNβ1 luciferase reporter vector (A-B) or ISRE (C-D) luciferase reporter vector along with an empty vector or a plasmid expressing the indicated protein. Doses of 300 ng, 100 ng, 30 ng and 3 ng per well of Nsp1 vectors and 300 ng, 200 ng and 100 ng per well of Nsp2 vectors were used. Thirty-two-hour post-transfection, cells were infected with SeV for 16h and then luciferase reporter activity was measured then standardized with a BCA protein dosage of the cell lysate. All experiments were done twice in triplicate and the compilation of the data is shown. Error bars indicate SD of experiments triplicates repetition. Statistical analyses were done by comparing corresponding condition with mock transfected control with a nonparametric one-way ANOVA with Dunn’s correction. *P<0,05, **P<0,01, ***P<0,001, ****P<0,0001. Relative protein expression levels of Nsp1 (E) and Nsp2 (F) with different transfection doses compared with SARS-CoV-2 infected A549-hACE2 cells. 24h post-seeding, A549-hACE2 cells were infected with SARS-CoV-2 and HEK293T were transfected as describe above. Cells were harvested 48h post-infection or transfection. Nsp1 and Nsp2 proteins were normalized with the correspondent Stain-Free blot lane (supplementary figure 1) and expressed as normalized levels (Norm. vol. x10^7^) below the blot.

### Nsp2 activates IFNβ1 production by activating NF-κB

As shown in the Fig. 5A the expression of Nsp2 without any stimulation activates the IFNβ1 promoter. To determine which regions of the IFNβ enhanceosome (Fig. 5B) were targeted by Nsp2, luciferase reporters containing either the positive regulatory domains I and III (PRD-I-III) (IRF3/7 responsive element) or the PRD-II (NF-κB-responsive element) were used (33). The luciferase reporters were co-transfected with the Nsp2 expression vector into HEK293T. We observed that Nsp2 activated the NF-κB binding domain of the IFNβ enhanceosome in a dose-dependent manner (Fig. 5C) while not affecting the PRD-I-III (Fig. 5D). These results indicated that Nsp2 activates the NF-κB pathway.

**Figure 5.**
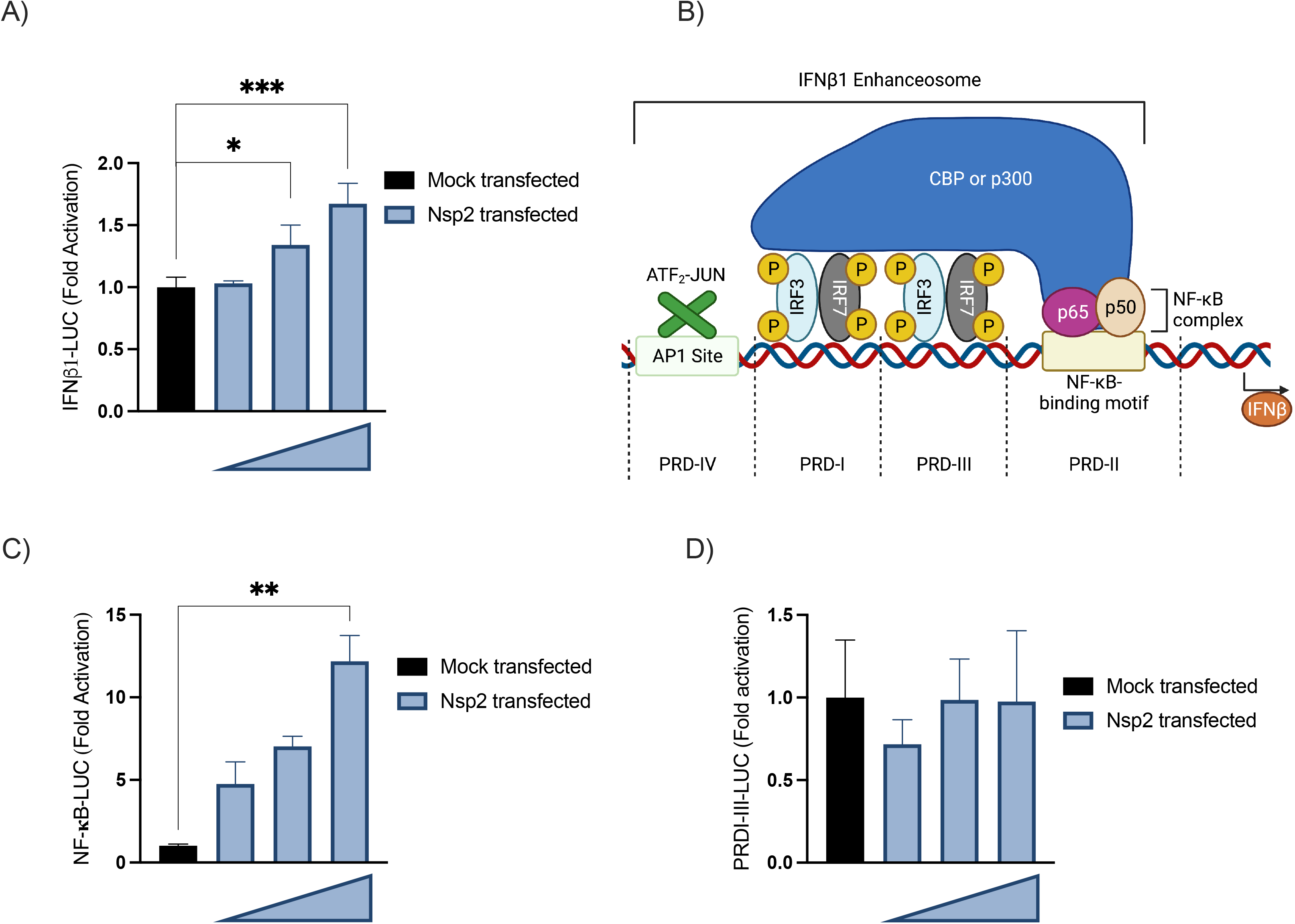
Characterization of IFNβ1 promoter activation by Nsp2. Effects of Nsp2 without stimulation on IFNβ1 promoter (A). Overview of the IFNβ1 promoter (B). Impact of Nsp2 expression on the NF-kB responsive elements, positive regulatory domain II (PRDII) (C) and IRF3 responsive elements, the positive regulatory domain I and III (PRDI-III) (D) of the IFNβ1 promoter. 300 ng, 200 ng and 100 ng per well of Nsp2 vectors were transfected along with the correspondent reporter and 48h post-transfection, the luciferase activity was measured then standardized as described for other reporter assays. All experiments were performed twice in triplicate, and one representative experimentation is shown. Error bars indicate SD of experiments triplicates. Statistical analyses were done by comparing mock transfected control with corresponding condition with a nonparametric one-way ANOVA with Dunn’s correction. *P<0,05, **P<0,01, ***P<0,001, ****P<0,0001.

### Nsp2 co-transfection fails to reduce the inhibitory effect of Nsp1

During infection, both Nsp1 and Nsp2 are produced simultaneously at an equimolar ratio. To determine whether both proteins might antagonize each other, IFNβ or ISRE luciferase reporter activation in response to SeV infection or IFNα was examined in co-transfection experiments. No significant difference between cells co-transfected with Nsp1 and Nsp2 vectors and cells singly transfected with Nsp1 vector alone was detected (Fig. 6A and B). In fact, cotransfection of Nsp2 failed to alter Nsp1’s ability to inhibit luciferase expression driven by the IFNβ and ISRE promoters. When transfected cells were analyzed for Nsp1 and Nsp2 expression, Nsp2 could be efficiently detected only in the absence of Nsp1, arguing that Nsp1 inhibited Nsp2 translation (Fig. 6C).

**Figure 6.**
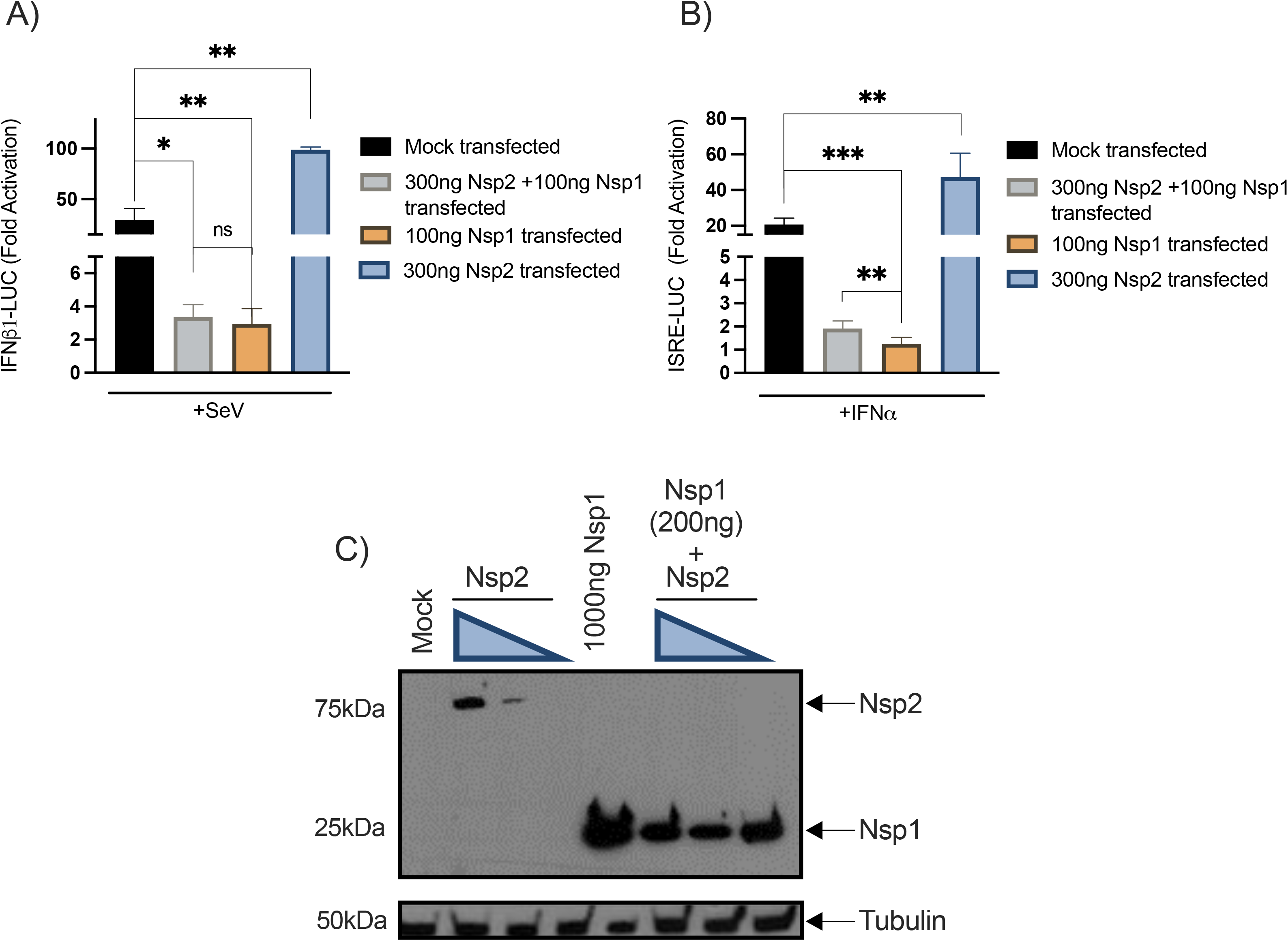
Effects of Nsp1 and Nsp2 cotransfection on SeV induced IFNβ1 promoter activation (A) and IFNα induced ISRE activation (B). HEK293T cells were seeded, transfected, and simulated according to the procedure described above and the luciferase activity was measured then standardized as described for other reporter assays. The Nsp1-Nsp2 cotransfection and Nsp1 transfection conditions were compared to the mock-transfected control using nonparametric one-way ANOVA with Dunn’s correction and Nsp2 conditions were compared using nonparametric one-way ANOVA. The Nsp1-Nsp2 cotransfection and Nsp1 transfection conditions were compared using nonparametric T-test. *P<0,05, **P<0,01, ***P<0,001, ****P<0,0001, ns: not significant. Protein expression of Nsp1 and Nsp2 in individual transfection and cotransfection (C). HEK293 were seeded in 6-well plate. 24h after, cells were transfected with control vector or Nsp1 expression vector or Nsp2 expression. 48h post-transfection, cells were lysed in SDS PAGE 2X buffer and detected by Western blot with an anti-FLAG (viral protein) and rabbit anti-tubulinβ (loading control).

### Nsp1-Nsp2 polyprotein coding vector succeeds to reduce Nsp1 inhibition on IFNβ pathways

To circumvent the fact that Nsp1 prevented the expression of Nsp2, we designed a vector expressing an Nsp1-P2A-Nsp2 polyprotein (schematized in Fig. 7A). Upon transfection of this polyprotein coding vector into HEK293T, the polyprotein along with Nsp1 and Nsp2 individual proteins were detected (Fig. 7B). The polyprotein vector was co-transfected with IFNβ1 or ISRE luciferase reporters into HEK293T cells. The results show that Nsp2 expression, enabled by the polyprotein vector, mitigated, at least partially, Nsp1 inhibition on the IFNβ1 reporters (Fig. 7C and D). Under the basal condition, a significant increase in IFNβ1 promoter activity was observed in the presence of Nsp1 and Nsp2 (Fig. 7C). In the presence of SeV, co-expression of Nsp1 and Nsp2 reduced the inhibitory effects of Nsp1 (Fig. 7D). However, co-expression of the two viral proteins did not modulate the Nsp1 inhibition on the ISRE reporter activity with mock and IFNα stimulated cells (Fig. 7E and F). These results indicate that Nsp2 partially antagonizes the inhibitory activity of Nsp1.

**Figure 7.**
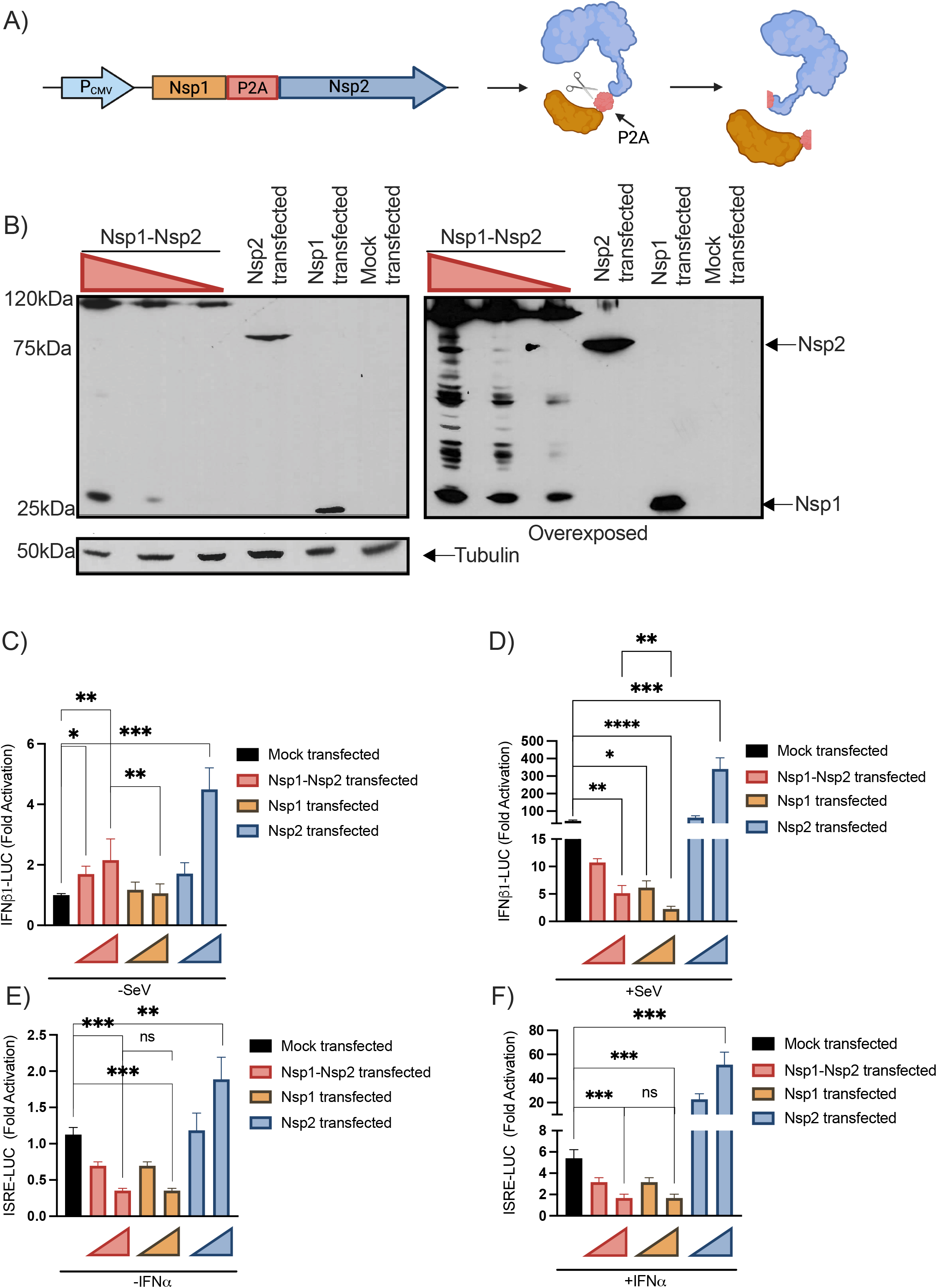
Effects of Nsp1-Nsp2 polyprotein on IFNβ and ISRE-luc promoter activation. Schematic representation of the Nsp1-P2A-Nsp2 encoding vector cleaved products (A). Nsp1 and Nsp2 expression upon transfection of the polyprotein coding vector. Vectors encoding Nsp1 and Nsp2 were used as controls (B). Effects of Nsp1-Nsp2 on IFNβ1-luc (C-D) and ISRE-luc (E-F) promoter activation. The Nsp1-Nsp2 and Nsp1 conditions were compared to the mock control using nonparametric one-way ANOVA with Dunn’s correction. The Nsp1-Nsp2 and Nsp1 conditions were compared together using nonparametric T-test. The Nsp2 transfection conditions were compared to the control using nonparametric one-way ANOVA with Dunn’s correction. *P<0,05, **P<0,01, ***P<0,001,****P<0,0001,ns: not significant.

### Nsp1 inhibits the IFNβ synthesis but does not affect the *IFNβ* gene transcription

To validate the result obtained using luciferase reporters, the effect of Nsp1, Nsp2, and Nsp1/Nsp2 co-expression on the *IFNB1*, *ISG15*, and *ISG56* mRNA and IFNβ1 protein production were measured. Our findings indicate that Nsp1 and Nsp2 do not affect *IFNB1*, *ISG15,* and *ISG56* genes transcription (Fig. 8A to D). In contrast, Nsp1 strongly inhibited the IFNβ1 production while the presence of co-expressed Nsp2 partially mitigated this effect (Fig. 8A). Overall, the data argue that Nsp1 shunts the IFN response by preventing the translation of mRNA, an effect partially antagonized by the Nsp2 protein.

**Figure 8.**
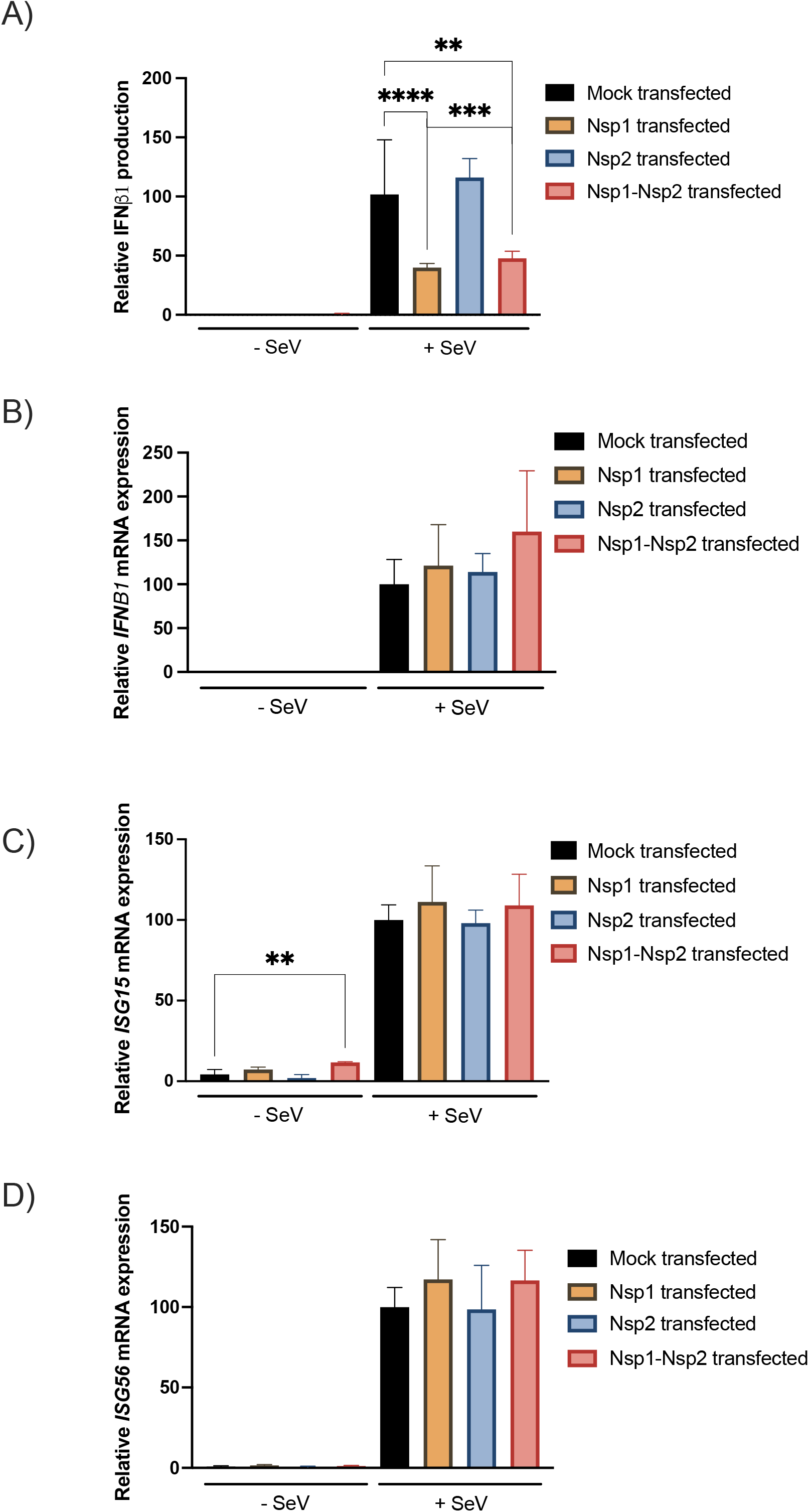
Effect of Nsp1, Nsp2 or Nsp1-Nsp2 polyprotein on *IFNB1* and *ISG* gene expression. Cells were transfected with expression vectors and infected or not with SeV as IFNβ1 inducer. Supernatants were collected and assayed for IFNβ1 production (A), RNA isolated and analyzed for *IFNB1* mRNA (B), *ISG15* mRNA (C) or *ISG56* mRNA (D) by RT-qPCR. All experiments were performed twice in triplicate. Results are expressed as activation percentage relative to the SeV infected control and error bars indicate SD of experiments triplicates repetition. Statistical analyses were done by comparing mock control with corresponding condition using nonparametric T-test. *P<0,05, **P<0,01, ***P<0,001, ****P<0,0001

## Discussion

In this study, we demonstrated that all SARS-CoV-2 strains tested induce robust *Ifnb1* gene transcription during *in vivo* infection of K18-hACE2 mice. Moreover, this *Ifnb1* gene expression was translated into detectable release of IFNβ1 in lung homogenates. *Ifnb1* gene expression is regulated by the coordinated actions of IRF3 and NK-κB, which are constitutively expressed in most cells (34, 35). Infected cells can therefore respond very rapidly to incoming viruses by inducing *Ifnb1* gene expression and IFNβ1 production even before viruses can deploy their anti-viral defense mechanisms. In the case of SARS-CoV-2, several viral proteins are reported to possess activities that antagonize the innate immune response such as type I IFN production. The most potent SARS-CoV-2 protein antagonizing the IFN response is the Nsp1 protein that induces a global shutdown of cellular mRNA translation (23, 24). The fact that several ISG, such as *Isg15, Igs56,* and *Mx1* are highly upregulated (Fig. 2B) during infection suggest that infected cells release sufficient IFNβ1 to induce the expression of genes associated with antiviral defense mechanisms. However, the establishment of an antiviral state is contingent on efficient ISG mRNA translation. To find out, we have examined *Irf7* gene expression and IFNα production. IRF7, constitutively expressed in plasmacytoid dendritic cells and B cells and induced in many other cell types by viral infections, is the main transcription factor responsible for the activation of IFNα promoters (12, 36). Plasmacytoid dendritic cells and B cells cell types do not appear to express ACE2 nor transmembrane serine protease 2 (TMPRSS2) (6), suggesting that they cannot be directly infected by SARS-CoV-2. In response to infection by all SARS-CoV-2 strains tested, *Irf7* is among the genes most highly expressed by all infected mice (Fig. 2B), suggesting that the recognition of infection by cellular sensors and downstream signaling molecules is functional. Differences at the level of pan *Ifna* gene expression and IFNα2/4 production were however observed between viral strains (Fig. 1C and D). Wuhan-infected mice had both significant pan *Ifna* gene induction and IFNα2/4 production relative to mock-infected and concordant with *Irf7* gene expression levels. In contrast, Beta- and Delta-infected mice had pan *Ifna* gene and IFNα2/4 levels that were equivalent to mock-infected mice, suggesting that the translation of IRF7 mRNAs is likely impaired. This would be consistent with the proposed role of Nsp1 (23, 24). However, the fact that the Nsp1 protein sequence is identical between Wuhan, Beta and Delta would argue that an alternative, yet to be identified mechanisms, can also affect IRF7 and/or *Ifna* genes expression. In that regard, the increased viral load RNA in Beta- and Delta-infected mice relative to Wuhan infected mice (Fig. 1A) might suggest that certain viral proteins are made at higher levels favoring greater immune evasion. IFNα inhibition in Beta-infected K18-hACE2 mice compared to the Wuhan strain was not observed in the work of Radvak & al. (37). In fact, a robust IFNα production was detected in lungs of Beta virus infected mice. This apparent discrepancy could possibly be explained by the lower virus inoculum (10^2^ TCID_50_) used in their study relative to ours (9×10^3^ TCID_50_). Additional differences with the study by Radvak and co-workers were also noted. For instance, the production of several cytokines in response to Beta infection, such as MIP1 α and β, were produced at high levels in this study.

Relative to type I IFNs and interleukins, CC and CXC chemokines were produced at high levels during infection by all SARS-CoV-2 strains in agreement with observations made in lungs of humans infected with SARS-CoV-2 and suffering from severe COVID-19 (38). In lungs of patients with severe COVID-19, CXCL8 (IL-8) and CXCL1 (GROα) were the predominant CXC chemokines. Mice do not encode *Cxcl8* gene, but do have CXCL1, which was produced at high levels (Fig. 1C and D). As in humans, CCL2 was the most prominent CC chemokines produced during infection (Fig. 1C and D). While CXCL1 and CXCL8 (human) are mainly involved in the neutrophil recruitment and activation, CCL2 is the main cytokine implicated in the monocyte recruitment as well as TH1 polarisation (39–41). Put together, concerted actions of these chemokines likely lead to a massive recruitment of leukocytes responsible of the acute respiratory distress syndrome (ARDS) observed in severe COVID-19 case (42).

Banerjee A & al. (27) recently reported that SARS-CoV-2 efficiently induced a type I IFN transcriptional response upon infection of pulmonary epithelial cells. Our work supports similar findings. In contrast, work by others (23, 24) clearly shows that this virus can also strongly inhibit the IFNβ1 protein expression. This apparent contradiction can be explained by the fact that certain studies measure RNA expression, while others evaluate protein synthesis. In fact, knowing that SARS-CoV-2 Nsp1 suppresses mRNA translation, the study of both mRNA and protein synthesis is necessary (23, 24). In that regard, our work confirms that infection of pulmonary epithelial cells by SARS-CoV-2 induced *IFN*β1 gene expression and even potentiated the response to IFN-inducing agents such as poly(I:C) (Fig. 3B to D). When IFNβ1 production in the supernatant was assessed however, partial inhibition in IFNβ1 production was measured only when infection was combinate with poly(I:C) stimulation (Fig. 3E). Considering that several other non-structural viral proteins are simultaneously generated with Nsp1 upon cleavage of the ORF1ab polyprotein, we surmised that at least one of them may partially antagonize the effects of Nsp1. Original to this work, we provided evidence that Nsp2, through activation of the NF-κB dampens the inhibitory effect of Nsp1 on IFNβ1 production. This effect of Nsp2 could be demonstrated when Nsp1-Nsp2 were generated from a common polyprotein alike the situation during viral infection (Fig. 7). However, when expressed individually, Nsp1 prevented the efficient expression of Nsp2 (Fig. 6C). Since SARS-CoV-2 infected cells do produce some IFNβ1 in response to infection suggest an incomplete blockage of mRNA translation by Nsp1 arguing that in addition of Nsp2, other viral factors may affect the activities of Nsp1. Our findings further highlight the potential caveats of studying viral proteins individually outside the context of infection as reported in several studies (25, 26, 43, 44). Nsp1 inhibition of IFNβ1-luc reporters has been shown by many, including this study. However, to our knowledge, this is the first report demonstrating that Nsp2 activates NF-κB (Fig. 5C). Nsp2-mediated activation of IFNβ1 and NF-κB reporters did not translate into increased IFNβ1 production. Nsp2 activation might be too small to affect IFNβ1 production in a measurable way but could play a role in the global immune response triggered by SARS-CoV-2 during infection.

While minimally inducing Type I IFN production in mice and *in vitro* model, early infection induces robust NF-κB activation driving the expression of chemokines like CXCL1,9,10,11 or CCL2,3,4,5 suggesting that virus skews the immune response toward an exaggerated inflammatory response rather than an antiviral response, as previously hypothesized (45). This overwhelming inflammatory response represents a major determinant of pathogenesis and morbidity observed during COVID-19. In support, the use of dexamethasone, a non-specific anti-inflammatory drug has proven effective in reducing mortality and length of hospital stay for patients with COVID-19 requiring oxygen supply (46).

In summary, the current study reveals that SARS-CoV-2 infection triggers vigorous expression of antiviral and inflammatory genes. However, in both mice and cell lines, IFN synthesis is sub-optimal, a consequence of the translational shutdown mediated by Nsp1. IFN shutdown in infected cells is however incomplete, in part due to the action of other viral proteins such as Nsp2 that partially antagonize the actions of Nsp1. As such, our work highlights the importance of studying viral protein functions in the context of infection. The use of recombinant mutant viruses will be helpful in delineating the synergistic/antagonizing functions of non-structural and accessory proteins during infection. In contrast to IFN, elevated inflammatory gene expression did translate into the production of high levels of several inflammatory chemokines, many of which are regulated by NF-κB. Considering our results demonstrating that Nsp2 activates the NF-kB pathway, Nsp2 should be considered as a potential contributor to the pathogenesis observed during SARS-CoV-2 infection.

## Acknowledgments

We thank the Laboratoire de Santé Publique du Québec for providing the Wuhan-like SARS-CoV-2 isolate used in this study. This study was supported by the Canadian Institutes for Health Research (CIHR) operating grant to the Coronavirus Variants Rapid Response Network (CoVaRR-Net) to LF, New Frontier research Funds (LF and EB), CIHR grants (VR3-172632, VS1-175516, VS1-175567) (LF and EB) and the CFI project “Seeking for innovative approaches in prevention and cure to COVID-19”. We thank the Centre de Recherche du CHU de Quebec sequencing platform for their excellent service. Émile Lacasse is the recipient of a fellowship from the Fondation du CHU de Québec. EB is recipient of an award from the Fonds de Recherche en Santé du Québec. Schematic representations of IFNβ1 and Nsp1-Nsp2 polyprotein vector (Fig. 5B and 7A) were created with BioRender.com.

---

**Supplementary Figure 1.** Uncropped blots (A-B) and Stain-Free blot (C-D) corresponding to the relative protein expression level of Nsp1 (A, C) and Nsp2 (B,D) within different transfection conditions compared with SARS-CoV-2 infected A549-hACE2 cells.

**Supplementary table 1.**
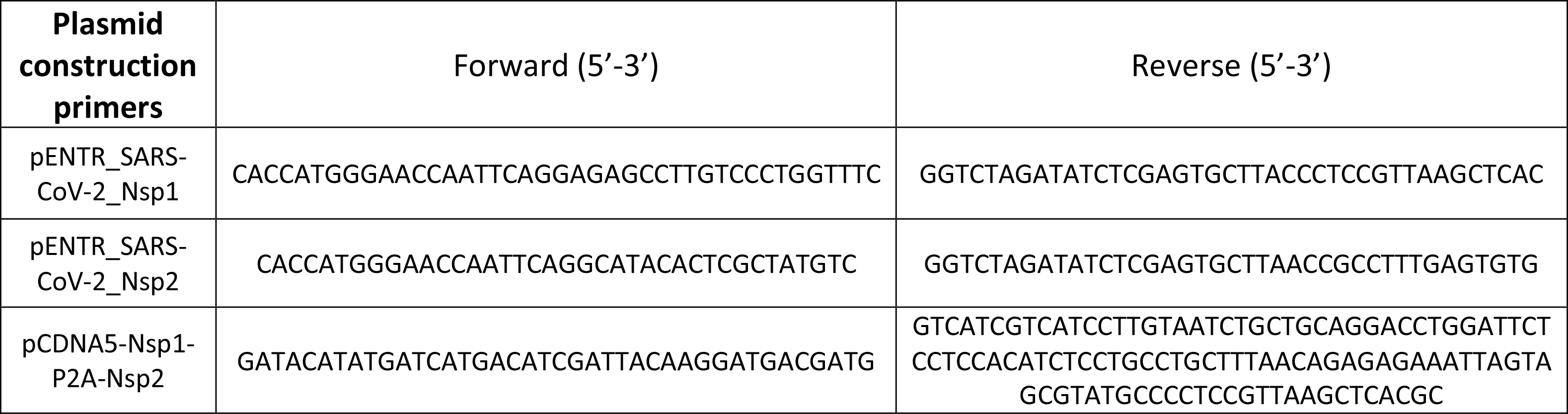
Primers used for plasmid construction.

**Supplementary table 2.**
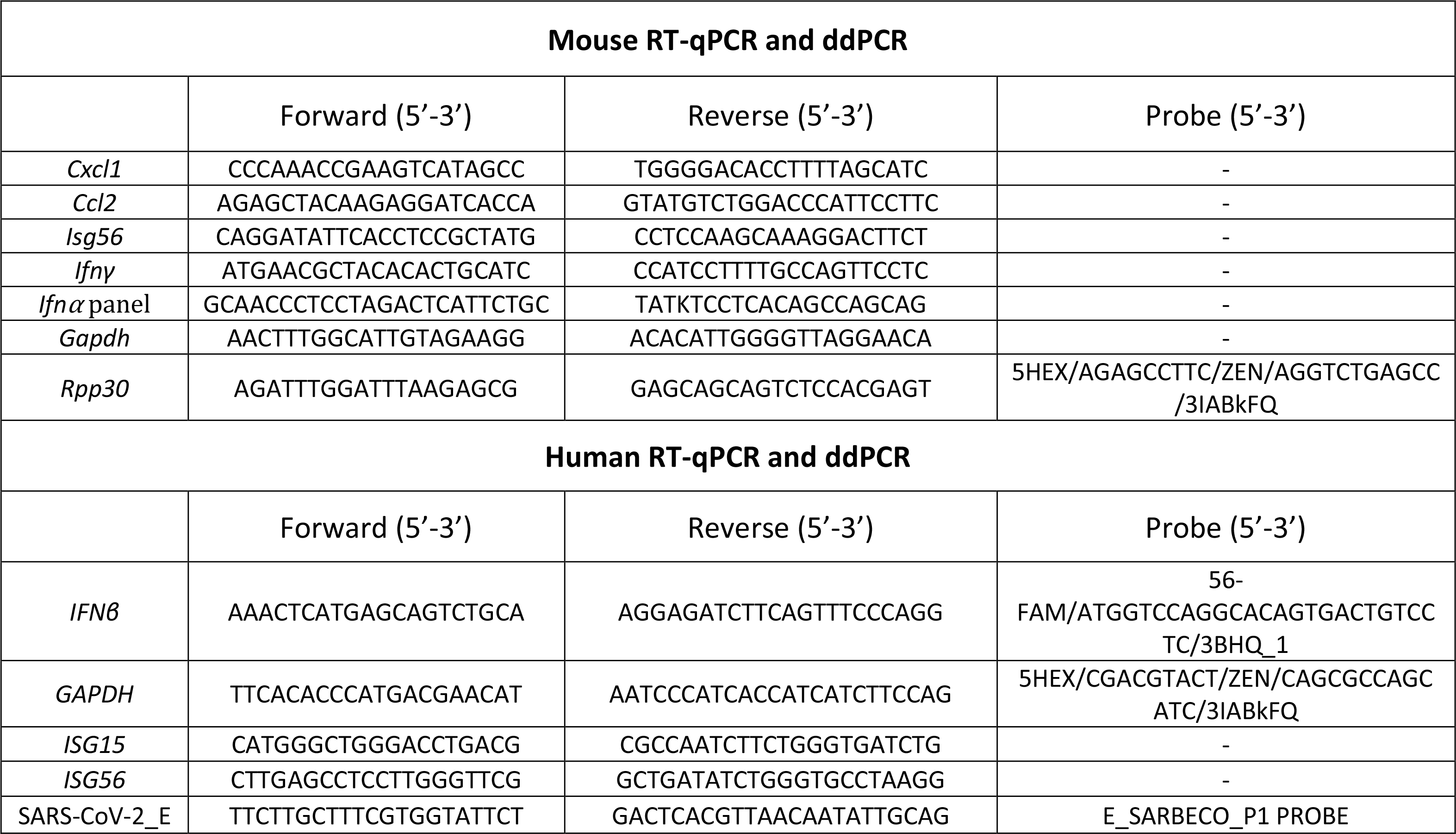
Primers and probes used for RT-qPCR and ddPCR experimentations.

**Supplementary table 3.** Genes included in RT^2^ profiler PCR Arrays Mouse Antiviral Response (PAMM-122ZR).

| Symbol | Description | Symbol | Description |
| --- | --- | --- | --- |
| Aim2 | Absent in melanoma 2 | Map2k1 | Mitogen-activated protein kinase kinase 1 |
| Atg12 | Autophagy-related 12 (yeast) | Map2k3 | Mitogen-activated protein kinase kinase 3 |
| Atg5 | Autophagy-related 5 (yeast) | Map3k1 | Mitogen-activated protein kinase kinase kinase 1 |
| Azi2 | 5-azacytidine induced gene 2 | Map3k7 | Mitogen-activated protein kinase kinase kinase 7 |
| Card9 | Caspase recruitment domain family, member 9 | Mapk1 | Mitogen-activated protein kinase 1 |
| Casp1 | Caspase 1 | Mapk14 | Mitogen-activated protein kinase 14 |
| Casp8 | Caspase 8 | Mapk3 | Mitogen-activated protein kinase 3 |
| Ccl3 | Chemokine (C-C motif) ligand 3 | Mapk8 | Mitogen-activated protein kinase 8 |
| Ccl4 | Chemokine (C-C motif) ligand 4 | Mavs | Mitochondrial antiviral signaling protein |
| Ccl5 | Chemokine (C-C motif) ligand 5 | Mefv | Mediterranean fever |
| Cd40 | CD40 antigen | Mx1 | Myxovirus (influenza virus) resistance 1 |
| Cd80 | CD80 antigen | Myd88 | Myeloid differentiation primary response gene 88 |
| Cd86 | CD86 antigen | Nfkb1 | Nuclear factor of kappa light polypeptide gene enhancer in B-cells 1, p105 |
| Chuk | Conserved helix-loop-helix ubiquitous kinase | Nfkbia | Nuclear factor of kappa light polypeptide gene enhancer in B-cells inhibitor, alpha |
| Cnpy3 | Canopy 3 homolog (zebrafish) | Nlrp3 | NLR family, pyrin domain containing 3 |
| Ctsb | Cathepsin B | Nod2 | Nucleotide-binding oligomerization domain containing 2 |
| Ctsl | Cathepsin L | Oas2 | 2'-5' oligoadenylate synthetase 2 |
| Ctss | Cathepsin S | Pin1 | Protein (peptidyl-prolyl cis/trans isomerase) NIMA-interacting 1 |
| Cxcl10 | Chemokine (C-X-C motif) ligand 10 | Pstpip1 | Proline-serine-threonine phosphatase-interacting protein 1 |
| Cxcl11 | Chemokine (C-X-C motif) ligand 11 | Pycard | PYD and CARD domain containing |
| Cxcl9 | Chemokine (C-X-C motif) ligand 9 | Rela | V-rel reticuloendotheliosis viral oncogene homolog A (avian) |
| Cyld | Cylindromatosis (turban tumor syndrome) | Ripk1 | Receptor (TNFRSF)-interacting serine-threonine kinase 1 |
| Dak | Dihydroxyacetone kinase 2 homolog (yeast) | Spp1 | Secreted phosphoprotein 1 |
| Ddx3x | DEAD/H (Asp-Glu-Ala-Asp/His) box polypeptide 3, X-linked | Stat1 | Signal transducer and activator of transcription 1 |
| Ddx58 | DEAD (Asp-Glu-Ala-Asp) box polypeptide 58 | Sugt1 | SGT1, suppressor of G2 allele of SKP1 (S. cerevisiae) |
| Dhx58 | DEXH (Asp-Glu-X-His) box polypeptide 58 | Tank | TRAF family member-associated Nf-kappa B activator |
| Fadd | Fas (TNFRSF6)-associated via death domain | Tbk1 | TANK-binding kinase 1 |
| Fos | FBJ osteosarcoma oncogene | Tbkbp1 | TBK1 binding protein 1 |
| Hsp90aa1 | Heat shock protein 90, alpha (cytosolic), class A member 1 | Ticam1 | Toll-like receptor adaptor molecule 1 |
| Ifih1 | Interferon induced with helicase C domain 1 | Tlr3 | Toll-like receptor 3 |
| Ifna2 | Interferon alpha 2 | Tlr7 | Toll-like receptor 7 |
| Ifnar1 | Interferon (alpha and beta) receptor 1 | Tlr8 | Toll-like receptor 8 |
| Ifnb1 | Interferon beta 1, fibroblast | Tlr9 | Toll-like receptor 9 |
| Ikbkb | Inhibitor of kappaB kinase beta | Tnf | Tumor necrosis factor |
| Il12a | Interleukin 12A | Tradd | TNFRSF1A-associated via death domain |
| Il12b | Interleukin 12B | Traf3 | Tnf receptor-associated factor 3 |
| Il15 | Interleukin 15 | Traf6 | Tnf receptor-associated factor 6 |
| Il18 | Interleukin 18 | Trim25 | Tripartite motif-containing 25 |
| Il1b | Interleukin 1 beta | Actb | Actin, beta |
| Il6 | Interleukin 6 | B2m | Beta-2 microglobulin |
| Irak1 | Interleukin-1 receptor-associated kinase 1 | Gapdh | Glyceraldehyde-3-phosphate dehydrogenase |
| Irf3 | Interferon regulatory factor 3 | Gusb | Glucuronidase, beta |
| Irf5 | Interferon regulatory factor 5 | Hsp90ab1 | Heat shock protein 90 alpha (cytosolic), class B member 1 |
| Irf7 | Interferon regulatory factor 7 | MGDC | Mouse Genomic DNA Contamination |
| Isg15 | ISG15 ubiquitin-like modifier | RTC | Reverse Transcription Control |
| Jun | Jun oncogene | PPC | Positive PCR Control |

